# A procedure for controlling the false discovery rate of *de novo* peptide sequencing

**DOI:** 10.1101/2025.09.12.675837

**Authors:** Justin Sanders, William Stafford Noble, Uri Keich

## Abstract

*De novo* sequencing is a powerful method for identifying peptides from mass spectrometry proteomics experiments without the use of a protein database. However, applications of *de novo* sequencing are currently severely limited by the lack of a reliable procedure for controlling the false discovery rate (FDR). Here, we introduce an FDR control procedure for the *de novo* setting which is at least as powerful as database search, give empirical evidence that it is statistically valid, and demonstrate its utility on a set of common *de novo* applications.

---

Recently, significant advances in *de novo* peptide sequencing have been made by us and others using deep learning models trained on massive datasets of labeled mass spectra [1]. These methods promise to enable the analysis of proteomics data in settings where a reference proteome is unavailable, which includes important applications to immunopeptidomics [2], environmental proteomics [3], antibody sequencing [4], venom sequencing [5] and paleoproteomics [6]. Additionally, when used in conjunction with database search, *de novo* sequencing can reveal discoveries typically missed by standard analysis pipelines, such as variant peptides [7, 8] and products of alternate translation [9]. However, a major roadblock limiting the widespread adoption of *de novo* tools is a lack of rigorous method for false discovery rate (FDR) control for *de novo* predictions. When peptides are assigned to acquired mass spectra via database search, target-decoy competition is the standard method for estimating FDR [10]. However, no analogous FDR-controlling procedure exists for the case of *de novo* peptide sequencing. This forces users to rely on only rough heuristics when applying *de novo* sequencing [3, 11–13] and prevents the assignment of any sort of statistical confidence in results.

In this work, we leverage the observation that, for essentially every application of *de novo* sequencing, a reference database containing at least a subset of relevant proteins or peptides is available. For example, when working with a species without a reference proteome, you can match some peptides to the nearest evolutionary homolog. When sequencing an antibody, you will have some peptides from the conserved region that match to the reference. More generally, it is rare that all peptides in a given sample are truly *de novo*, even if the discoveries of interest are those not in the reference. Accordingly, we propose a method for controlling the FDR for *de novo* sequencing results which is applicable specifically in the setting where some, but not all, peptides in the sample come from a known sequence database.

While prior work has explored the concept of FDR control for *de novo* sequencing in this setting, current approaches suffer from a number of major limitations and thus have not reached general adoption. When performing their FDR control procedure, PostNovo [14] considers only spectra that were identified by database search, but the selected score threshold is then applied to all predictions. The spectra identified by database search are not an unbiased sample, because they are likely to be enriched for high-quality spectra. Novoboard relies on the generation of perfectly convincing decoy spectra, which is a major challenge due to both the expressiveness and non-interpretability of the deep learning models used in modern *de novo* sequencing algorithms. Finally, and most importantly, all existing methods control the FDR on the combined set of *de novo* and database discoveries. [14–16] This approach is problematic because typically users are running *de novo* sequencing because they are interested specifically in making *de novo* discoveries. If the FDR is controlled on the combined list, then the FDR among just the set of non-reference *de novo* discoveries may be much higher. Incorrect *de novo* peptide predictions are almost certain to not be in the reference, whereas correct predictions may or may not match the reference database. Indeed, this problem of FDR control in a subset has already been pointed out in the literature in the database search context [17, 18].

Recognizing these limitations, we propose a two-stage procedure for FDR control, in which we subject a given set of spectra to both database search and *de novo* sequencing. We first use well established decoy-based methods to control the FDR among the database search results. Thereafter, we propose a novel procedure to control the FDR among *de novo* discoveries from the residual spectra which remain unidentified by database search. We present an argument that, subject to two intuitively reasonable assumptions, the procedure controls the FDR among this set of *de novo* discoveries (See Methods).

Our proposed two-stage procedure offers two important benefits. First, the method is always at least as powerful, in a statistical sense, as database search. This overcomes the low sensitivity of *de novo* sequencing, which is a major limitation hindering more widespread adoption. Second, the method is agnostic with respect to the database search or *de novo* sequencing method. This means that, like target-decoy competition, our procedure can be added on top of any existing analysis workflow, provided some weak assumptions on properties of the *de novo* algorithm.

Given a collection of spectra from a proteomics experiment, our procedure begins by running both a database search algorithm and a *de novo* algorithm on each spectrum (Figure 1A). Spectra identified at the desired FDR threshold by database search are then removed, and the corresponding discoveries are included in the final list. For the remaining spectra, our procedure aims to make discoveries among the set of non-reference (unmatched) peptides predicted by the *de novo* algorithm. We call these predictions “external peptides,” and they consist of a mixture of both correct and incorrect predictions. The proportion *π*_0_ of incorrect predictions in the mixture is unknown and hence must be estimated by our procedure. The procedure works by using agreements between *de novo* predictions and database search to estimate the distribution of *de novo* scores assigned to correct discoveries.

**Figure 1.**
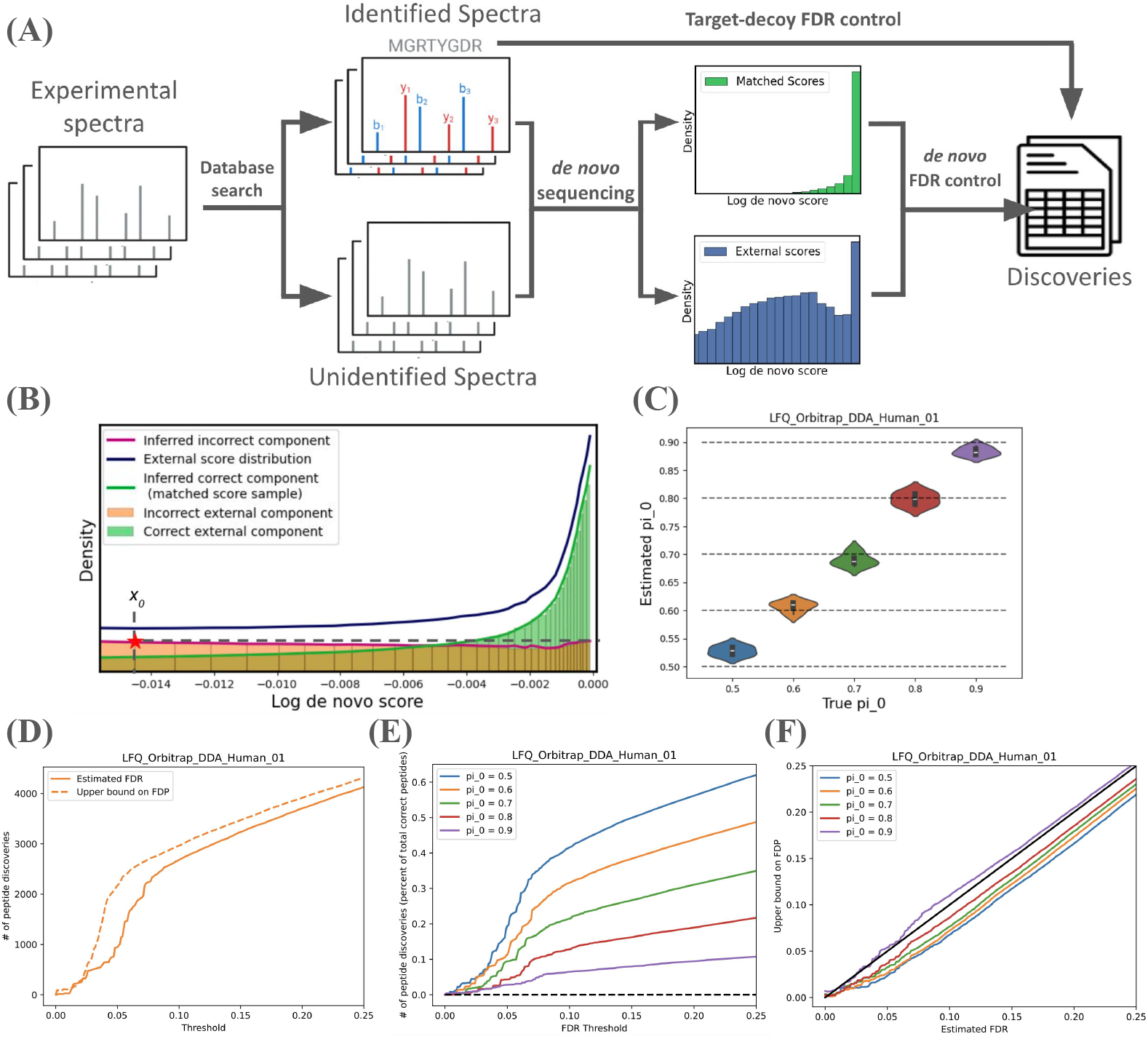
The proposed FDR control procedure. (A) A schematic overview of the procedure. Experimental spectra are identified using database search and *de novo* sequencing, where the latter are split to those that match the reference database and those that are external to the database. The FDR is estimated based one the assumption that the external *de novo* scores represent a mixture of incorrect and correct matches where the latter share the same distribution of the *de novo* scores of the matched predictions. (B) A plot comparing the probability-density functions of the correct and incorrect components (green and orange histograms, respectively) of the external score distribution to the densities inferred by our procedure (green and magenta line). The estimated antimode 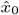 is indicated with a vertical dashed line, and the density at 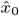, which is used for the stopping criterion of the procedure, is indicated with a horizontal dashed line. (C) Violin plots showing the distribution of inferred *π*_0_ across bootstraps in each experiment. The estimated *π*_0_ is both accurate and precise. (D) A comparison of the number of peptide discoveries made by our procedure at each FDR threshold to the number discovered at the same FDP threshold calculated based on the ground truth peptide labels. (E) FDR curves showing the average number of *de novo* discoveries made at each FDR threshold in across a range of true values of the mixing parameter *π*_0_ . The y-axis is normalized to the proportion of the total number of ground truth peptides present in the experiment. (F) Comparison of the average estimated FDR to an upper bound on the true FDP across a range of experimental conditions. The FDR of our procedure is well controlled across a range of different mixing parameters *π*_0_

At a high level, our approach offers a twist on the canonical approach to FDR control where decoys are used to model the score distribution of incorrect discoveries. In our approach, we instead model the score distribution of correct discoveries by using matches to reference peptides discovered by database search. Just as in the target-decoy strategy, we then use this estimated correct score distribution to separate the mixture distribution of external scores into its correct and incorrect components.

Our procedure is based on the observation that, given an empirical estimate of the correct score distribution, if we knew the mixing parameter *π*_0_ then we could directly read off the FDR at a given score threshold based on the relative cumulative likelihood under each distribution. Thus, given an estimate of the distribution of the correct scores, controlling the FDR on the set of external peptides boils down to accurately inferring *π*_0_. Unfortunately, *a priori* this is an underconstrained problem. To make it tractable, we introduce two assumptions on the shape of the score distributions.

Specifically, we assume, first, that the distribution of *de novo* scores assigned to correct predictions is the same for both matched (reference) and external predictions, and that the probability density function (PDF) of this distribution is monotone increasing. Second, we assume that the distribution of the scores of incorrect *de novo* predictions is unimodal with its mode *x*_*m*_ *<* 0 (0 is the maximal possible score). Furthermore, in order for our procedure to work, the external scores mixture PDF needs to have a rightmost local minimum (antimode) *x*_0_ ∈ (*x*_*m*_, 0). Our experience is that this is often the case but at this point while we are still working on devising a complete test for that the user needs to visually confirm this by examining the histogram of their external scores. If the given data is not consistent with the existence of *x*_0_, as will typically happen if the mixture PDF is increasing, then our procedure should not be used.

Our core procedure begins with finding an estimate 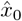 of the antimode *x*_0_. We next estimate *π*_0_ by finding the largest *ρ* ∈ (0, 1) such that after subtracting from the mixture PDF the matched score PDF scaled by *ρ*, the remaining estimated true null PDF does not exceed its estimated value at 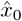 for any point 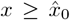 (Figure 1B). To smooth out the effects of stochasticity in selecting 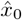 and estimation of *π*_0_, we repeat the above procedure across many bootstrapped samples of the original data and return the average results.

To evaluate our procedure, we set up a simulated *de novo* sequencing setting where the ground truth label for *de novo* predictions is known. To do this, we start with a standard mass spectrometry experiment from a species with a well characterized proteome, in which we expect there to be very few true *de novo* discoveries to be made. We then hold out a portion of the proteome, effectively creating a set of peptides which can only be discovered *de novo* but which we know to be correct. We then run our procedure, obtaining a set of external discoveries at a given FDR threshold. We estimate the false discovery proportion (FDP) on this set by counting how many of the predicted peptides are from the held-out portion of the reference proteome, which are counted as true discoveries, versus how many are not in the reference, which are counted as false discoveries. In practice there may be a small number of other *de novo* peptides present in the sample, e.g., from unaccounted-for contaminants. This makes the evaluation a conservative upper bound, meaning that if the FDR is less than the estimated FDP then the FDR is likely controlled, but if it is greater, then either the FDR control is too permissive or the bound on the FDP is too loose. However, if we assume that true *de novo* peptides are relatively rare in this data, then the resulting upper bound on the FDP should be tight. By varying the proportion of the reference proteome held out, and thus the number of true de novo discoveries to be made in the experiment, we can simulate various settings with different values of the mixing parameter *π*_0_.

Evaluating our procedure using this experimental setup using the Tide search engine and the Casanovo *de novo* sequencer [19, 20], we find that the resulting FDR appears to be well-controlled albeit somewhat conservative across a wide range of values of *π*_0_ (Figure 1C). As *π*_0_ approaches 1, the estimated FDR appears more liberal, and at times even surpasses the upper bound on the FDP. This is not too unexpected, as we expect the upper bound to grow looser as *π*_0_ increases — a small number of real *de novo* peptides from contaminants will have a larger effect when the number of possible true discoveries is small. Further testing our procedure on an additional eight experiments in four species from three different instrument platforms, we find that the results look highly consistent, indicating that our two assumptions consistently hold in practice (Supplementary Figures 3-5). Additionally, we show that the method is applicable to results from multiple de novo sequencing algorithms (Supplementary Figure 6).

To demonstrate the utility of our procedure in a real world application, we analyze data collected from bone fragments of the extinct cave bear *Ursus deningeri* [21]. Due to the lack of reference proteomes for extinct species, paleoproteomics relies on searching data against the reference proteome for the most closely related sequenced extant species, and thus can benefit from *de novo* sequencing to identify peptides that do not appear in the reference. Here, as in the original analysis by Johnson *et al*., we run our procedure using the grizzly bear (*Ursus arctos horribilis*) reference proteome, which is the closest living relative to the cave bear. We identify 197 database peptides and 35 *de novo* peptides at a 1% FDR. Among the *de novo* discoveries, we find many exact matches to other closely related bear species, such as giant panda (*Ailuropoda melanoleuca*) and black bear (*Ursus americanus*) (Figure 2B). This offers strong evidence into which of the alleles observed in extant bear species were carried by *Ursus deningeri*, and demonstrates how FDR controlled *de novo* sequencing enables new statistically rigorous insights into the proteome of this extinct species.

**Figure 2.**
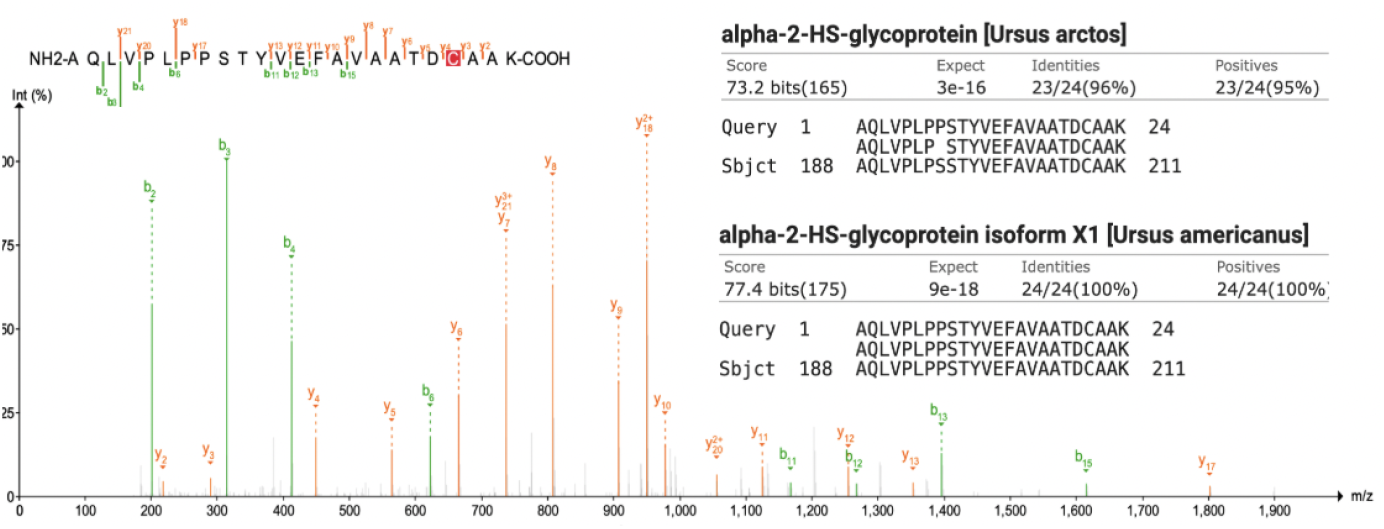
FDR control enables analysis of downstream tasks. (An example spectrum for a *de novo* peptide discovered at 1% FDR from the cave bear dataset. The top scoring alignments to the *U. Arctos* and *U. Americanus* reference proteomes are shown in inset.

In conclusion, we present a new method for separately controlling the FDR among both database and *de novo* discoveries from the same analysis. Notably, the procedure is strictly more powerful than database search alone, meaning that *de novo* sequencing can be added to any workflow without incurring a loss in sensitivity. These results address many of the current limitations hindering the more general adoption of *de novo* sequencing methods. We have implemented our procedure in an open source Python package, available at https://github.com/Noble-Lab/glissade, that is designed to work with any search engine and *de novo* sequencing algorithm.

## Methods

### Formal description of the procedure

#### Problem setup

Our procedure takes as input two sets of search results for the same run — one from a standard database search controlled at a pre-selected FDR threshold (we used 1%) and one from a *de novo* sequencing algorithm — along with the reference proteome for the sample which was used during database search. The *de novo* sequencing results are split into two sets, those where the *de novo* peptide and database search result for a given scan agree, which we will call a matched PSM, and those where the *de novo* peptide is not in the reference and the scan was not identified by database search, dubbed an external PSM. Predictions where *de novo* sequencing and database search disagree, along with predictions which are in the reference but not identified by database search, are discarded. This avoids a double-dipping problem of attempting to make discoveries on the same set of database peptides twice, which would inflate the number of false discoveries. Results are then aggregated to the peptide level by taking the top scoring PSM for each peptide, yielding a set of matched peptides and a set of external peptides. If not already representing log-likelihoods, the scores assigned to these two sets of peptides are transformed to the interval [™∞, 1] by first standardizing to the interval [0, 1] and then taking the log-transform. The procedure below aims to control the FDR among discoveries from the set of external peptides.

Let *S*_*m*_ be the set of scores assigned to the matched peptides, and let *S*_*e*_ be the set of scores assigned to the external peptides. We expect *S*_*e*_ to be a mixture distribution containing two components, those scores assigned to correct predictions and those assigned to incorrect predictions. Our goal is to separate the the mixture into its two components. To make this de-convolution possible, we we make the following two assumptions:

**Assumption 1** *The matched scores and the correct component of the external scores are independently drawn from the same distribution which has a monotone increasing PDF*.

**Assumption 2** *The distribution of the incorrect scores has a unimodal PDF with an internal mode x*_*m*_ ∈ (−∞, 0).

#### Estimating *x*_0_

As mentioned in the introduction, our core procedure is designed for the case where there exists a rightmost antimode *x*_0_. We then find an estimate of *x*_0_ by iteratively shrinking an interval *I* to identify the first “significant” local minimum of the estimated mixture density as we move away from the maximum of 0. This process starts with 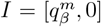, where 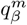 is the *β* quantile of the *N*_*m*_ matched scores. *β* is a tunable parameter of our procedure; here we used *β* = 0.2 for all results. Having initialized the interval *I*, we iteratively update it by the following steps:

1. Test whether the left third of *I* or the middle third of *I* are significantly enriched relative to the entire interval (each of these is a binomial test with *p* = 1*/*3 and the corresponding number of scores that fall within the relevant 1/3 of the interval versus the full intervals). If either of these tests is significant at level *γ* (we used *γ* = 0.05) then move the left end point of I forward by min{*n*_*s*_, |*I*|*/*3} points (we used *n*_*s*_ = 50) and start again with the new *I*.
2. Otherwise, check whether the right third of *I* is significantly enriched relative to the entire interval and if so move the right end point of *I* backward min{*n*_*s*_, |*I*|*/*3} points and go back to (1);
3. Otherwise, check if |*I*| *< n*_*s*_ and if so set 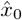 to the middle of *I* and terminate the procedure;
4. Otherwise, move both the left end point of *I* toward the center of the interval by min{*n*_*s*_, |*I*|*/*3}*/*2 and go back to (1).

#### Inferring *π*_0_

Having found an estimate for the antimode *x*_0_ we next aim to estimate the mixing parameter of the external score distribution *π*_0_. We do this by iteratively resampling with replacement from the set of matched scores *S*_*m*_. Under Assumption 1, this matched sample has the same distribution as the correct component of the external scores. Thus, by “subtracting” this increasing sample away from the distribution of external scores we simulate a sample from the mixture distribution with a decreasing proportion of correct scores. We terminate this sampling process when the estimated PDF of the remaining mixture becomes negative or that it seems to satisfy Assumption 2. In the latter case we obtain a generally conservative estimate of *π*_0_ (because Assumption 2 should hold for the correct *π*_0_).

More specifically, we begin with associating with each external score *s*_*i*_ a window *W*_*i*_ that contains the *n*_*w*_ points that are closest to it. The parameter *n*_*w*_ is chosen in a data driven manner as a fraction of the total number of scores greater than 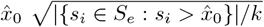, where *k* is a hyperparameter for the procedure (we used *k* = 5 for all results here).

We next sequentially draw with replacement from the matched score distribution *S*_*m*_. Let *m*_*j*_ for *j* = 1, …, *ℓ* be the *ℓ* hitherto drawn matched scores, and for each 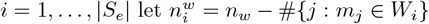. That is, 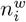 is the number of external scores in the window *W*_*i*_ minus the number of the drawn matched scores that happen to fall within the same window. At each step we check if Assumption 2 approximately holds by examining whether the estimated density in each window to the right of 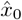 is not larger than at 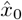. Specifically, with 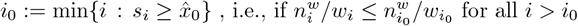. If this holds, or if there exists a window with an updated 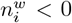, indicating it has negative density, we stop, where in the latter case we also subtract 1 from *ℓ* because the density should not be negative.

With the sampling process complete we view the sampled matched scores *m*_1_, …, *m*_*ℓ*_ as proxies for the correct scores among the external scores and hence we estimate the FDR at score threshold *s*_*i*_ as

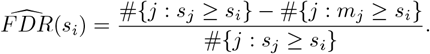

#### Bootstrapping the procedure

To reduce the effect of sampling variance on parameter inference and FDR estimation, we repeat the above procedure across *B* bootstraps. Within each bootstrap, we resample with replacement the external scores *S*_*e*_ before running the procedure to get a single FDR estimate for each score threshold 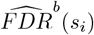 from that bootstrap. Finally, we report the average FDR across all bootstraps at each threshold:

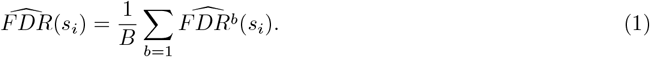

For all results here we use *B* = 100.

#### Consistency tests

As mentioned, the user needs to visually inspect the histogram of the external scores to verify that it is reasonable to assume *x*_0_ exists. We are working on developing a complete test for that, but in the meantime we already implemented the following test guarding against the mixture PDF being monotone decreasing on [*x*_*d*_, 0] for some *x*_*d*_ *<* 0. Specifically, we check whether the estimated density is increasing across the last *X* windows of the external scores. If that is not the case, we skip the above core of our procedure so no *de novo* discoveries are reported.

In addition, when our secondary stopping criterion in the above resampling process (there exists a window with an updated 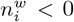)holds, we employ one more test to add some confidence in the validity of our assumptions. Specifically, we apply Kolmogorov-Smirnov to test whether the external scores that are greater or equal to the center of that triggering window and the matched scores come from the same distribution. If that test is rejected at the 0.05 level our procedure again reports no *de novo* discoveries.

### Validity of the FDR control

Our simulations do not reveal any substantial flaw in our procedure’s FDR control. Moreover, we expect our procedure to be generally asymptotically valid if our assumptions hold. That said, there could be pathological cases where the above procedure for estimating *x*_0_ might fail to do that in a reasonably accurate manner. In updates to this work we will consider alternative methods that will guarantee asymptotic convergence to *x*_0_ as the sample size grows.

### Experimental setup

#### Database search

Each dataset was searched against the appropriate species fasta using the Tide search engine (version XXX) with XXX. Data was searched with a tryptic digest allowing for one missed cleavage, an isotope error of 1 or 2, and up to three modifications. The modification list used was the same set of modifications considered by Casanovo (oxidation of methionine, deamidation of asparagine and glutamine, and N-terminal acetylation, carbamylation, and ammonia loss).

For the human data in this study, we use the UniProt Homo sapiens reference proteome UP000005640, downloaded on 15 February 2024. For mouse, we use the Mus musculus reference UP000000589, downloaded on 29 April 2024. For yeast, we use the UniProt Saccharomyces cerevisiae reference UP000002311, downloaded on 5 August 2024. For *E. coli*, we use the UniProt Escherichia coli reference UP000000625, downloaded on 2 February 2025. All four references were appended with a list of common protein contaminants [22].

#### *De novo* sequencing

Unless otherwise specified, *de novo* sequencing was performed using the Casanovo search engine (version 3.3.1) with precursor mass tolerance set to 1,000,000 to turn off the precursor filter, and all other options set to default.

For the DeepNovo results, the data was searched with a fork of DeepNovo version 0.0.1 downloaded from github on 4-2-25. By default, DeepNovo only reports scores to two significant figures, leading to a discrete score distribution which greatly undermined the power of our procedure. To circumvent this, we modified the DeepNovo source to report scores to full precision. All other aspects of the algorithm were left identical to version 0.0.1.

#### Evaluating the FDP

To evaluate the validity of our FDR control procedure, we set up an experiment where the ground truth labels of *de novo* discoveries is known. We start by running both database search and *de novo* sequencing on a sample with a well characterized proteome, yielding our lists of matched scores *S*_*m*_ and external scores *S*_*e*_. We then hold out a portion of the matched scores *S*_*c*_ ⊂ *S*_*m*_ and mix them into our set of external scores. This simulates the setting where the reference proteome is incomplete, and some true peptides may be identified only by *de novo* sequencing. By varying the number of correct peptides mixed in, we can vary the true *π*_0_ of the mixture distribution. We can obtain an upper bound on the FDP of our procedure by counting how many of the discoveries made at a specific score threshold come from this set of known correct sequences which we mixed in:

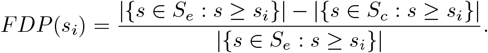

If we assume that there are no true foreign peptides in the data (i.e., peptides that are in the sample but not in the database), then this bound on the FDP will be perfectly tight. In practice, this bound is likely to be somewhat loose due to the presence of sequence variants and unconsidered contaminants. However, as long as we assume that this set is significantly smaller than the number of native peptides (i.e., peptides that are in the sample and in the database), then this upper bound will be tight. Additionally, as an upper bound on the FDP, it is sufficient for showing the validity of our procedure without any assumptions on the tightness of the bound, though it is unable to concretely determine whether our procedure is liberal or if the bound is loose.

To properly demonstrate that our procedure is working we need to show that for a given sample of correct scores, the FDP is well controlled in expectation across samples of incorrect scores from the null. To this end, we perform our evaluation experiments by running 100 iterations of our procedure across different re-samplings of the external scores *S*_*e*_ for a fixed set of correct scores *S*_*c*_, computing for each iteration both the estimated FDR using our procedure and the ground truth FDP as described above. We asses the validity of the estimated FDR by plotting the average FDP across iterations at each FDR threshold.

### Applications

#### Cave bear analysis

The raw data from the single cave bear bone fragment run, along with the *Ursus arctos horribilis* reference fasta, was downloaded from the PRIDE data repository PXD015083 [21]. Data was searched with the Tide search engine with hydroxyl-proline, hydroxyl-lysine and oxidized methionine included as variable modifications and deamidation of asparagine and glutamine into aspartic acid and glutamic acid set as fixed modifications. We ran our procedure on the data with default paramaters and accepted discoveries at an estimated FDR of 0.05. These peptide discoveries where then searched with BLAST [23] against all sequences from the Ursidae family (NCBI taxid:9632) the nr database to asses homology to other extant bear species.

## Data Availability

The MS/MS datasets used in this study were all downloaded from the pride repositiories PXD028735, PXD066485, and PXD001468 [24–26].

