## Supplementary figures for "A procedure for controlling the false discovery rate of *de novo* peptide sequencing"

### Supplementary information

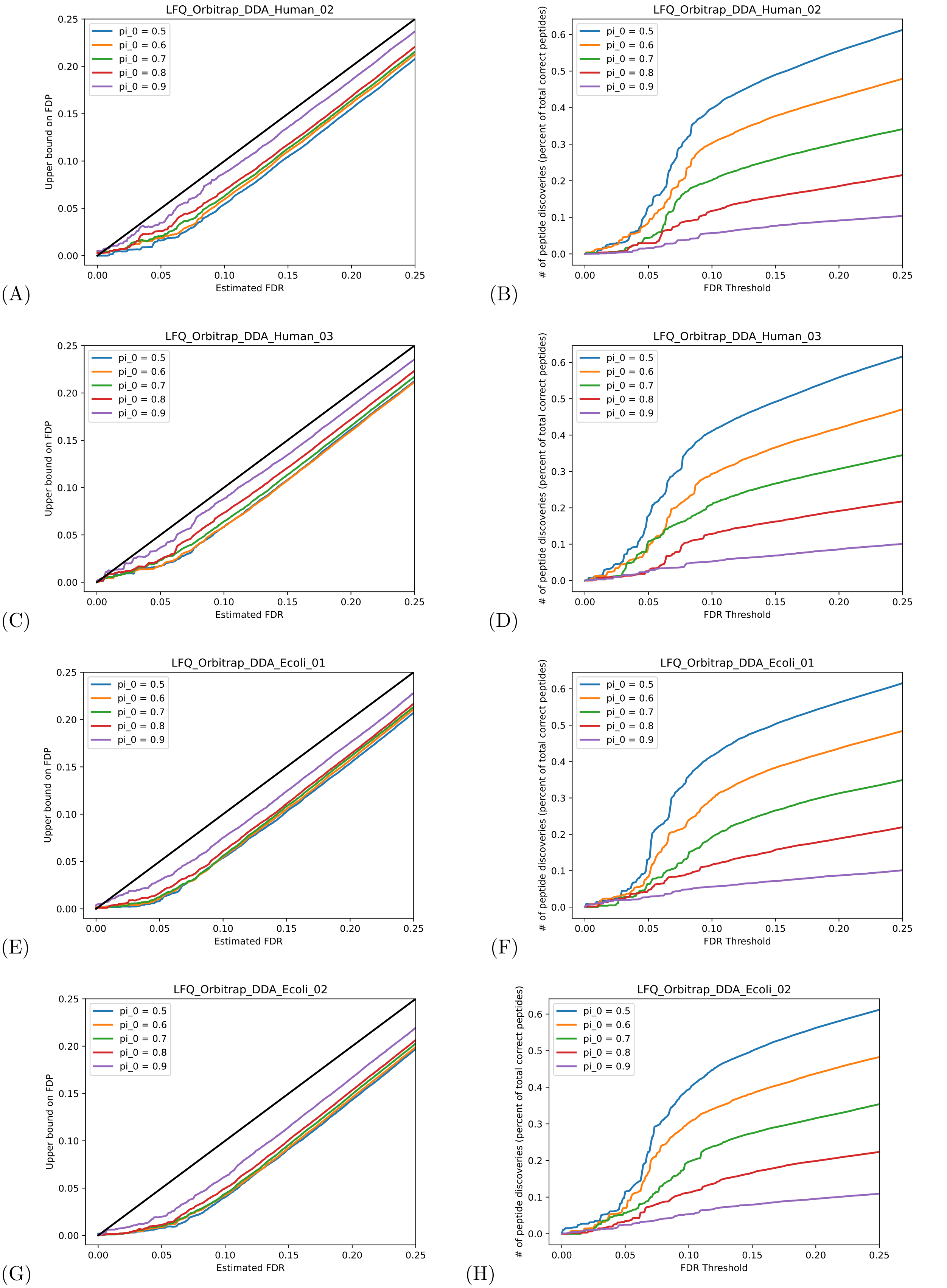


Figure 3: **Evaluation on additional datasets**. (A-H) Plots showing the calibration and power of our procedure on each mass spectrometry run from PXD028735 [24]

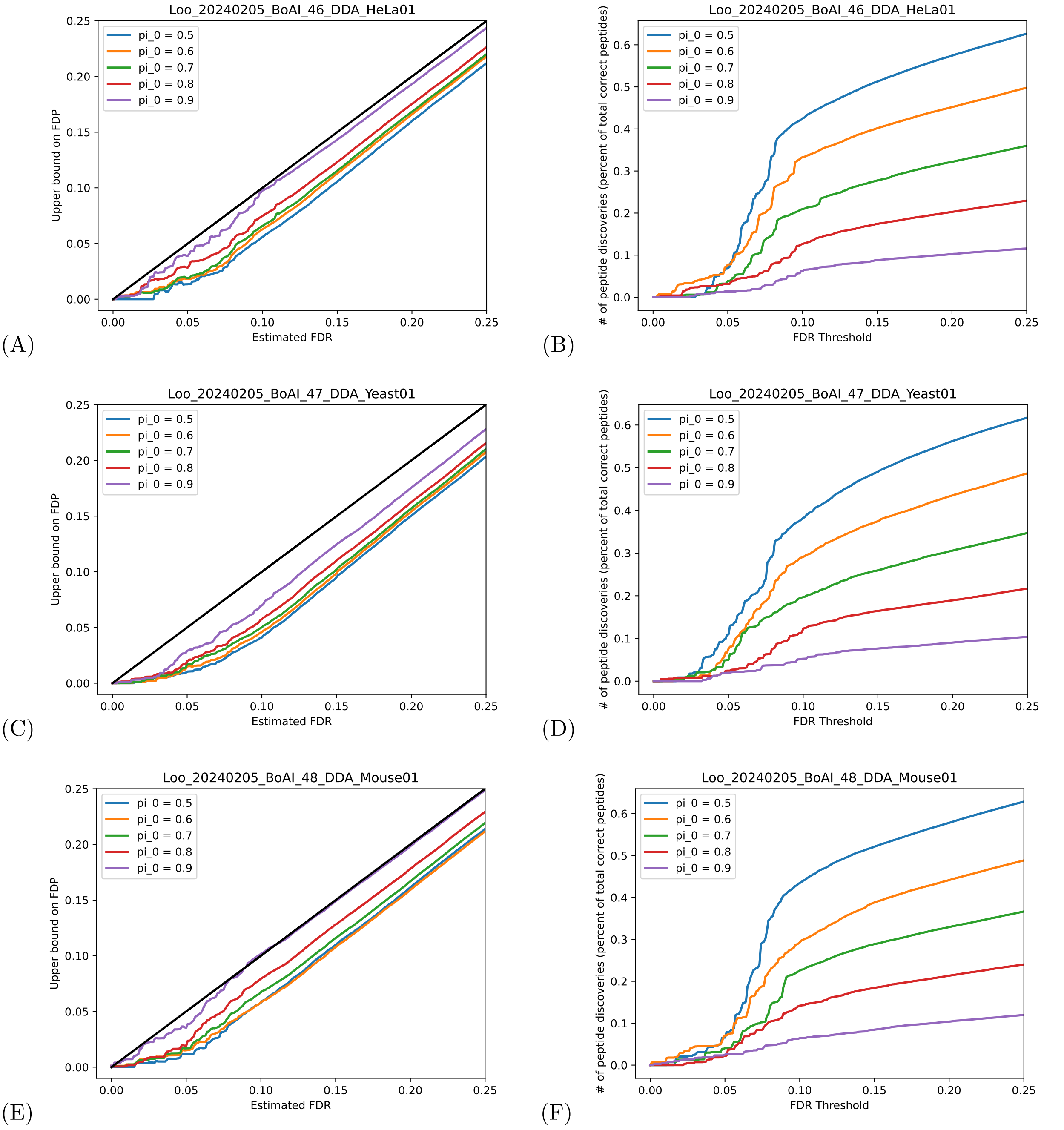


Figure 4: **Evaluation on additional datasets**. (A-F) Plots showing the calibration and power of our procedure on each mass spectrometry run from PXD066485 [25]

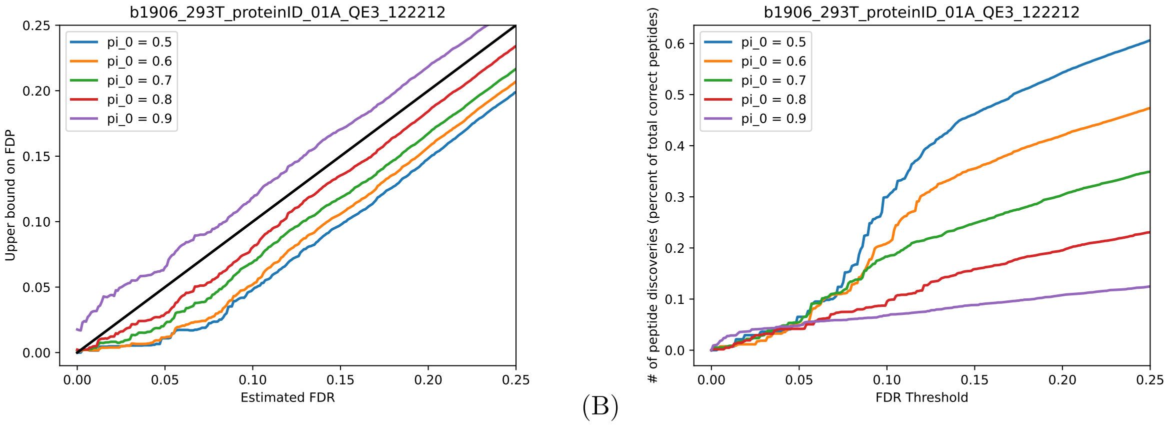


(A) (B)

Figure 5: **Evaluation on additional datasets**. Plots showing the calibration (A) and power (B) of our procedure on each mass spectrometry run from PXD001468 [26]

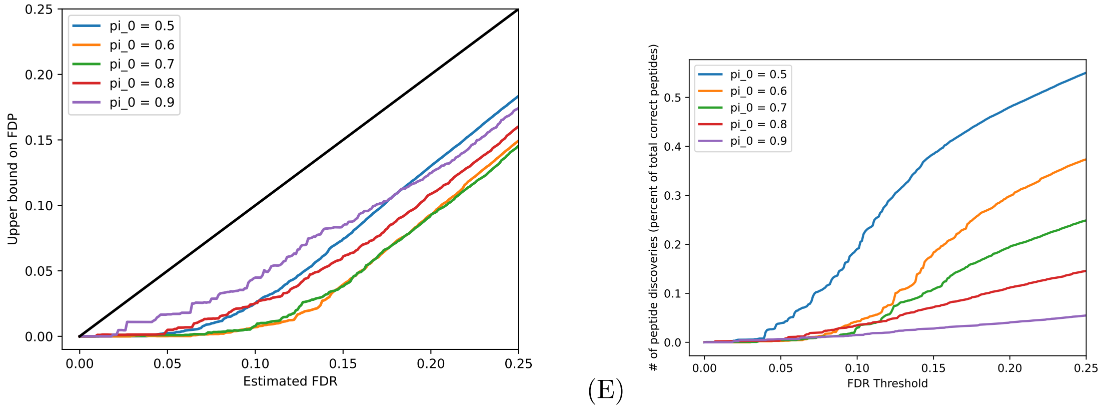


(F) (E)

Figure 6: **Evaluation with DeepNovo scores**. Plots showing the calibration (A) and power (B) of our procedure when applied to results from the *de novo* sequencing algorithm DeepNovo [27]
